# Artificial cells with all-aqueous droplet-in-droplet structures for spatially separated transcription and translation

**DOI:** 10.1101/2024.06.11.598395

**Authors:** Kanji Tomohara, Yoshihiro Minagawa, Hiroyuki Noji

## Abstract

The design of functional artificial cells involves compartmentalizing biochemical processes to mimic cellular organization. To emulate the complex chemical systems in biological cells, it is necessary to incorporate an increasing number of cellular functions into single compartments. Artificial organelles that spatially segregate reactions inside artificial cells will be beneficial in this context by rectifying biochemical pathways. In our study, we developed artificial cells featuring all-aqueous droplet-in-droplet structures that separate transcription and translation processes, mimicking the nucleus and cytosol in eukaryotic cells. This droplet-in-droplet architecture utilizes intrinsically disordered protein (IDP) to form coacervate droplets for the inner compartments, and aqueous two-phase systems (ATPS) for the outer compartments, with the outer interfaces stabilized by colloidal emulsifiers. The inner droplet was designed to enrich DNA and RNA polymerase to conduct transcription, which was coupled to translation at the outer droplet, realizing the cascade reaction mediated by mRNA. We also demonstrate that these processes proceed independently within each artificial cell compartment, maintaining the correspondence between genotype and phenotype. The modular configuration of these artificial organelles could be extended to other enzymatic reactions. Coupled with the ease of manufacturing these artificial cells, which only requires simple agitation in an all-aqueous mixture, this approach provides a practical and accessible tool for exploring complex systems of artificial organelles within large ensembles of artificial cells.

## Introduction

The bottom-up artificial cells have been studied in the field of synthetic biology, where cellular functions are reconstituted in cell-sized compartments. This research aims to identify the minimal composition of living systems and chemically distinguish them from non-living ones. It also offers an industrial platform for producing beneficial biomolecules while avoiding cellular toxicity and incorporating non-canonical elements.

Molecules for representative cellular functions, such as transcription-translation^1^, DNA replication^2–5^, and cellular division^6^, have typically been introduced to a wide range of compartments. Ideal compartments should be biocompatible to facilitate biological reactions internally and have the capability of self-organization for growth and division^7^. One prime example is a liposome, where the lipid bilayer surrounds the aqueous droplet akin to modern cells. Liposome has been extensively studied as an artificial cell compartment together with other membranous compartments.

On the other hand, membrane-less compartments like coacervates have been proposed to be a possible candidate for protocells before the advent of lipids^8,9^. These compartments are increasingly getting attention as artificial cell reactors because of their unique features of molecular permeability, selective molecular enrichment, and crowded interiors^10^.

These membrane-less compartments are formed through liquid-liquid phase separation (LLPS), which can be classified into two modes: segregative LLPS and associative LLPS. Among the segregative LLPS systems, one of the most well-studied involves a pair of neutral polymers, i.e., Dextran and PEG (polyethylene glycol), which form an ATPS (aqueous two-phase system)^11,12^. Both Dextran-in-PEG and PEG-in-Dextran droplets can be formed depending on the concentration of both polymers, and it has been shown that DNA, RNA, and most proteinous components for cell-free protein expression are enriched in Dextran-in-PEG droplets^13–16^, enabling protein expression inside. Meanwhile, associative LLPS gives coacervates formed through multivalent intermolecular interaction. It has been actively reported recently that low-complexity protein regions form these condensates inside cells and act as membrane-less organelles to control biochemical pathways spatiotemporally in a stimuli-responsive manner^17^.

The concept of organelles is also starting to be applied to artificial cells^18–20^. These artificial organelles in the artificial cell will offer chemically isolated space for specific reactions, thus enabling spatial arrangement of modular functions and higher-order organization of complex chemical systems closer to the framework of living organisms. Enzymatic cascade reactions have been demonstrated with compartment-in-compartment systems^21^. For example, hydrogels with DNA being chemically immobilized are encapsulated in compartments to allow transcription, followed by translation to express protein^22,23^. Alternatively, self-organizing membranous droplets also serve as the inner compartment, where this inner membrane can be engineered to permit the transfer of small molecules. These molecules can then be used as substrates for the enzymatic reactions in the outer compartments^21,24^.

Here, we show that the droplet-in-droplet structure can be formed in an all-in-water mixture by the combination of coacervate and ATPS and transcription and translation can be respectively conducted in the inner droplet and the outer droplet. The droplet-in-droplet structure can be spontaneously formed by mixing three polymers: IDP (intrinsically disordered protein) forming coacervate as the inner droplet and Dextran and PEG forming ATPS as a Dextran-in-PEG outer droplet. The interface between Dextran and PEG was stabilized by proteinous colloids to suppress coalescence, thus maintaining the individuality of the artificial cell compartments. The inner IDP droplet acted as a field for transcription by selective recruitment of template DNA and RNA polymerase. Moreover, thanks to the membrane-less nature of the inner droplet, transcribed mRNA can diffuse out to the outer Dex droplet and be translated to express proteins, realizing the cascade reaction. This system not only features the spatial hierarchical structure of the nucleus and the cytosol in eukaryotic cells but also mimics their respective role to enable compartmentalized transcription and translation.

## Result

### Formation and stabilization of droplet-in-droplet structure by LLPS

First, we formed a droplet-in-droplet structure by mixing three polymers: IDP (intrinsically disordered protein), Dex (Dextran), and PEG (polyethylene glycol). This IDP is a construct of RGG-BFP-RGG, where RGG is an intrinsically disordered region derived from LAF-1 in *Caenorhabditis elegans* and responsible for creating membrane-less droplets as coacervates^25,26^, and BFP (mTagBFP2, blue fluorescent protein) enables observation of droplets with fluorescence microscopy. While Dex and PEG are known to form segregative LLPS droplets as ATPS (aqueous two-phase system), we found that the addition of IDP to this system resulted in droplet-in-droplet structures. As shown in Figure 1a and Figure 1b, the IDP droplet nested inside the Dex droplet, surrounded by a continuous PEG phase.

**Figure 1.**
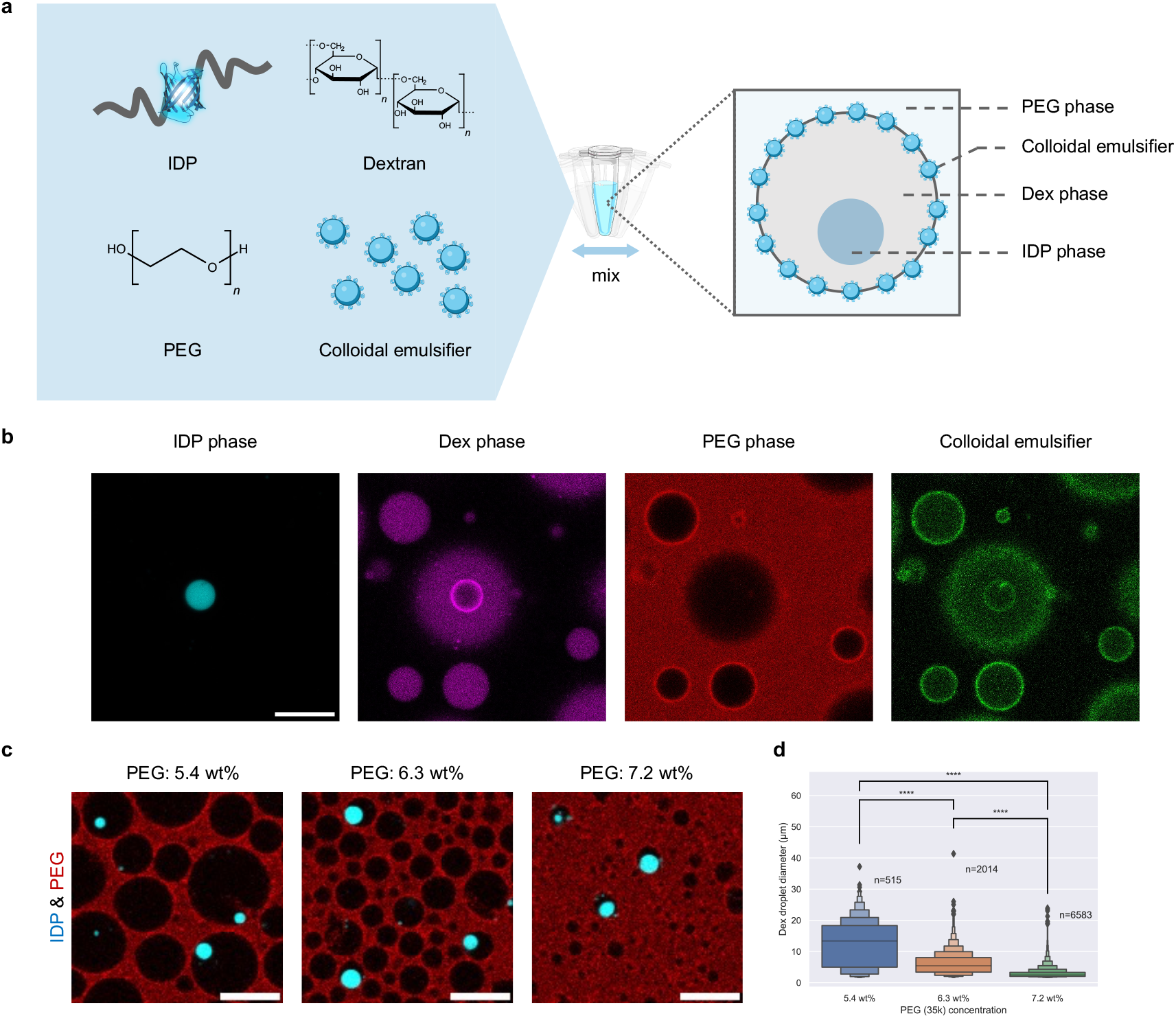
Surface-stabilized droplet-in-droplet structure by polymer LLPS. **a** Schematic of the preparation process of the surface-stabilized droplet-in-droplet structure. Dextran (Dex) and PEG undergo segregative LLPS to form Dex and PEG phases, whose interface is stabilized by PEGylated proteinous colloids as emulsifiers. Inside the droplet of the Dex phase, intrinsically disordered protein (IDP) forms a coacervate droplet. **b** Confocal microscopy images of each phase and emulsifiers. IDP is genetically fused with BFP for fluorescence observation. The Dex phase and the PEG phase are fluorescently labeled by TRITC-Dex and Atto647N-PEG, respectively. Proteinous colloidal emulsifiers are labeled by SYPRO orange. The scale bar represents 10 μm. **c**,**d** Dex droplets’ size tunability by altering the concentration of PEG. Scale bars indicate 20 μm. Significance was assessed using a two-tailed Welch’s t-test: NS (p > 0.05), * (p < 0.05), ** (p < 0.01), *** (p < 0.001), **** (p < 0.0001).

However, Dex droplets is prone to coalescence, which is problematic in artificial cell compartments; it is crucial to maintain a correlation between each genotype and phenotype. While the interface between oil and water can be simply stabilized by the addition of amphiphilic molecules, this is not the case for the interface of Dex and PEG phases because both are aqueous phases^27^. Hence, Dex-PEG interface is typically stabilized as Pickering emulsion instead, where microparticles surround the dispersion phase and prevent coalescence between dispersion phases^28,29^. In this study, we utilized protein-based colloidal microgels as emulsifiers prepared by heat-denaturation of BLG (β-lactoglobulin). BLG colloids are known to stabilize PEG-in-Dex droplets at neutral pH, but they do not stabilize Dex-in-PEG droplets^30^. Given the principle that effective emulsifiers should favor the continuous phase over the dispersed phase^31^, we PEGylated the BLG colloids. This modification stabilized Dex-in-PEG droplets for over four hours (Supplementary Figure 1).

While the total volume fractions of the Dex and PEG phases in an ATPS can be controlled by adjusting their concentrations, the incorporation of BLG colloids, which help prevent droplet fusion, allowed us to control the size of each dispersed Dex droplet. As shown in Figure 1c, by setting the PEG concentration to 6.3 wt% in the presence of BLG colloids, we successfully achieved Dex droplets within the biologically relevant range of 5 to 20 μm. This tunability enables the preparation of artificial cells at the scale of natural cell structures.

### Localization of transcription in IDP droplet

Next, we aimed to conduct transcription at the inner IDP droplets (Figure 2a). To achieve this, we localized essential transcription components—RNA polymerase (RNAP) and template DNA—within the IDP droplet. To recruit proteins, including enzymes, to the IDP droplets, Schuster *et al*. utilized the interaction between peptide tags SYNZIP1 (SZ1) and SYNZIP2 (SZ2), which were fused to IDP and the proteins of interest (POI), respectively^25,32^. We implemented this approach by attaching the SZ2 tag to T7 RNA polymerase (T7RNAP) and the SZ1 tag to IDP. Enrichment of T7RNAP in the IDP droplet was assessed bay labeling either SZ2-tagged T7RNAP or non-tagged T7RNAP with fluorescein (FL), followed by measuring the fluorescence in both the IDP and Dex phases. While T7RNAP without the SZ2 tag already demonstrated a 14-fold enrichment factor, defined as the ratio of fluorescence in the IDP droplet to that in the Dex droplet, the SZ2-tagged T7RNAP exhibited a 190-fold enrichment factor, as illustrated in Figure 2b, 2c and Supplementary Figure 2. Similarly, we attached an SZ2 tag to the lac repressor (LacR), enabling the DNA coding the lac operator (lacO) sequence to interact with this modified SZ2-LacR. This modulation, combined with the SZ1-SZ2 and LacR-lacO interactions, facilitated the recruitment of the lacO DNA to the IDP droplet. This was verified by labeling lacO DNA with FL and analyzing its distribution in the droplet-in-droplet structure both with and without SZ2-LacR (Figure 2d, 2e). Consequently, the enrichment factor of lacO DNA increased from 7.9 to 45 with the addition of SZ2-LacR. Since the present study aims to perform translation in addition to transcription, all these observations were conducted in the presence of cell-free translation components (i.e., PURE frex 2.0 ΔT7RNAP).

**Figure 2.**
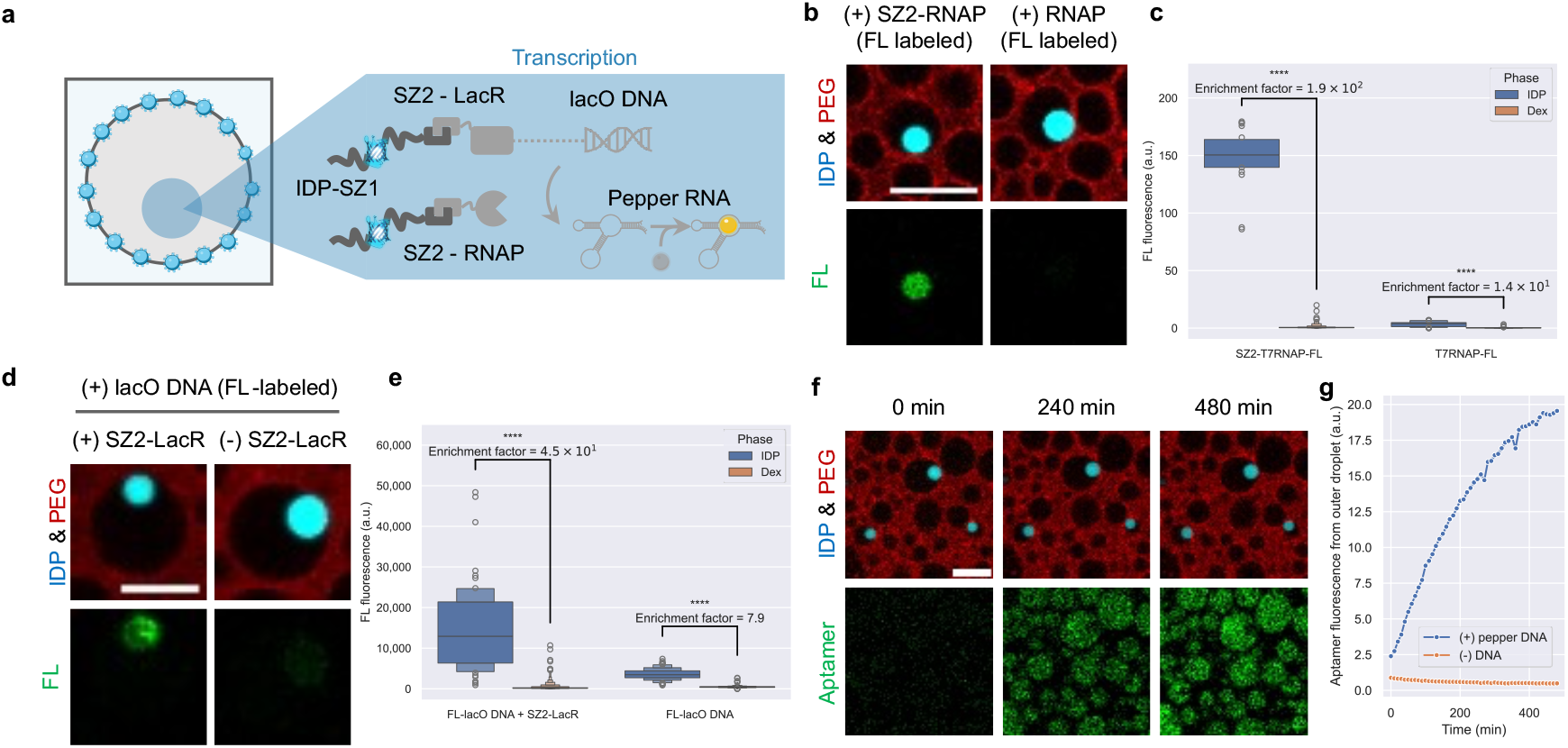
Transcription in the inner IDP droplet. **a** Schematic of transcription in the inner IDP droplet. Enzymes (RNAP) and repressors (LacR) are both designed to bind with IDP through SZ1-SZ2 peptide interactions. LacR binds to the lacO sequence placed outside the region to be transcribed, thus recruiting template DNA to the IDP droplet. Transcribed RNA contains pepper aptamer for fluorescence observation. **b**,**c** Enrichment of SZ2-tagged RNAP confirmed by fluorescence label. **d**,**e** Enrichment of fluorescence-labeled lacO-containing DNA in the presence or absence of SZ2-LacR. **f** Time-lapse images of transcription monitored by aptamer fluorescence. **g** Line plot of aptamer fluorescence from the outer Dex droplet with or without pepper DNA. Error bands represent 2SE. All scale bars represent 10 μm. Significance was assessed using a two-tailed Welch’s t-test: NS (p > 0.05), * (p < 0.05), ** (p < 0.01), *** (p < 0.001), **** (p < 0.0001).

To conduct time-lapse observation of RNA synthesis under a microscope, we employed the Pepper fluorescent RNA aptamer along with its specific activating dye, HBC^33^. The template DNA included the F30 scaffold and eight tandem repeats of the Pepper RNA genes, all under the control of the T7 promoter, with two lacO sequences positioned upstream of the transcription region. This setup allowed us to monitor the transcription activity by the increase in aptamer fluorescence (Figure 2f). As shown in Figure 2g, the aptamer fluorescence in the outer Dex droplet continuously increased over 480 minutes, in contrast to the condition where Pepper DNA was not added. Further analysis revealed that both the inner IDP droplet and the outer Dex droplet exhibited comparable levels of fluorescence signal after 480 minutes of expression (Supplementary Figure 3), showing that RNA diffused out and would be available for subsequent translation in the outer Dex droplet when mRNA was transcribed.

**Figure 3.**
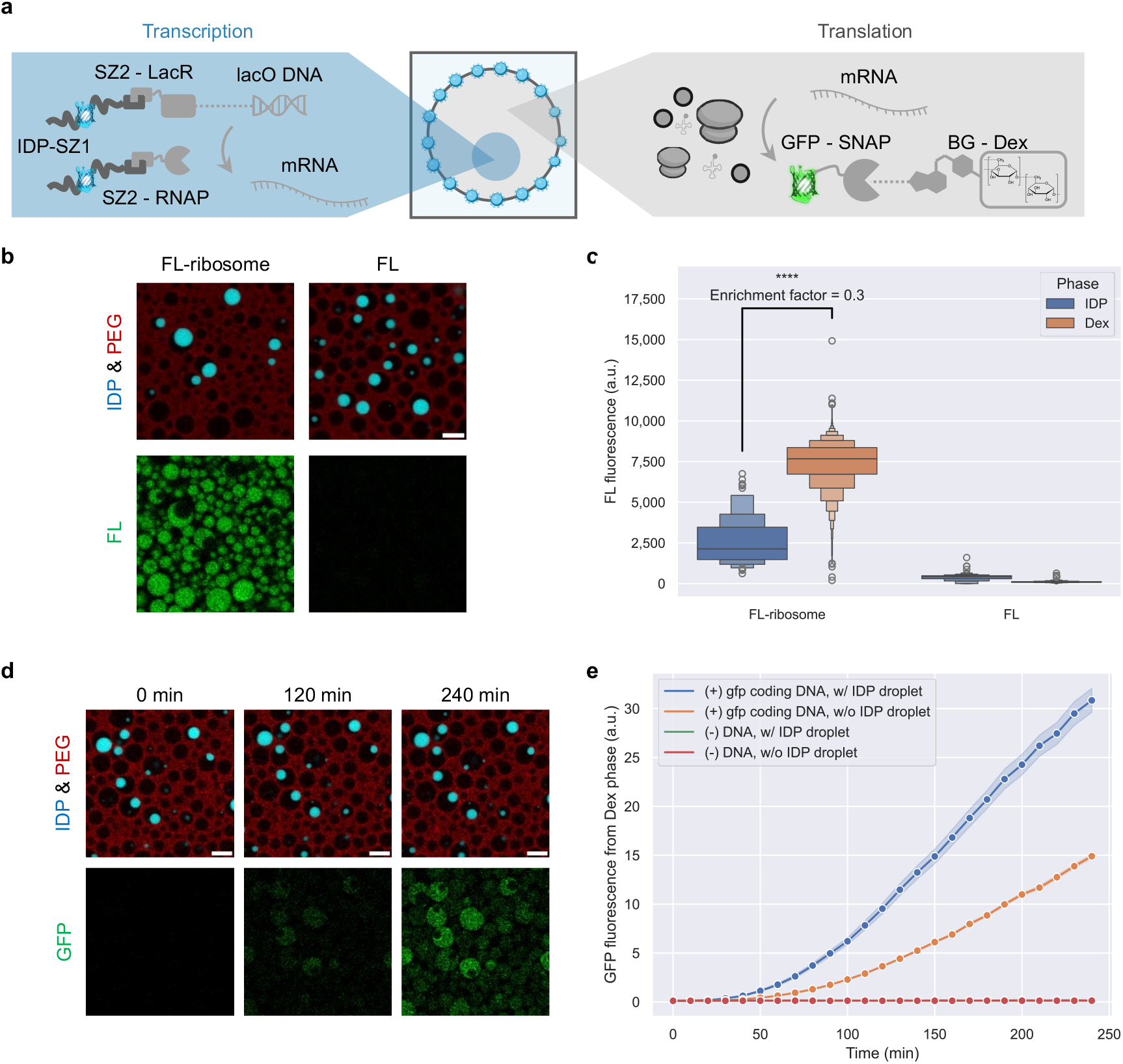
Translation at the outer Dex droplet coupled with transcription at the inner IDP droplet. **a** Schematic. Translation at the outer Dex droplet from mRNA synthesized in the inner droplet. By expressing GFP tagged with SNAP-tag in the presence of benzyl guanine-Dextran (BG-Dex), GFP can be fixed to the Dex phase. **b**,**c** The location of ribosomes confirmed by dye-labeled ribosomes (0.1 μM) with confocal microscopy. **d** Time-lapse images of translation. **e** GFP expression from Dex droplets with or without inner IDP droplets were compared. Error bands represent 2SE. All scale bars represent 10 μm. Significance was assessed using a two-tailed Welch’s t-test: NS (p > 0.05), * (p < 0.05), ** (p < 0.01), *** (p < 0.001), **** (p < 0.0001).

Alongside microscopic observations, we performed bench-top experiments with conventional test tubes to ensure that transcription proceeds in the IDP phase rather than the Dex phase. The IDP-Dex phase system was centrifuged and macroscopically separated into a supernatant Dex phase and a sedimented IDP phase in the presence of PURE frex 2.0 ΔT7RNAP, SZ2-T7RNAP, SZ2-LacR and HBC (Supplementary Figure 4a). Since the volume of IDP phase was too small (about 1/10^3^ of the whole volume) to extract directly, we sampled the supernatant Dex phase and compared it to the whole mixture sampled before centrifugation (IDP and Dex phases). Monitoring transcription activities with aptamer fluorescence, it was revealed that the RNA yield at 170 minutes was two-fold higher in the whole mixture compared to the supernatant Dex phase (Supplementary Figure 4d). Given the minute volume fraction of the IDP phase, this result indicates that transcription preferentially took place in the IDP phase.

**Figure 4.**
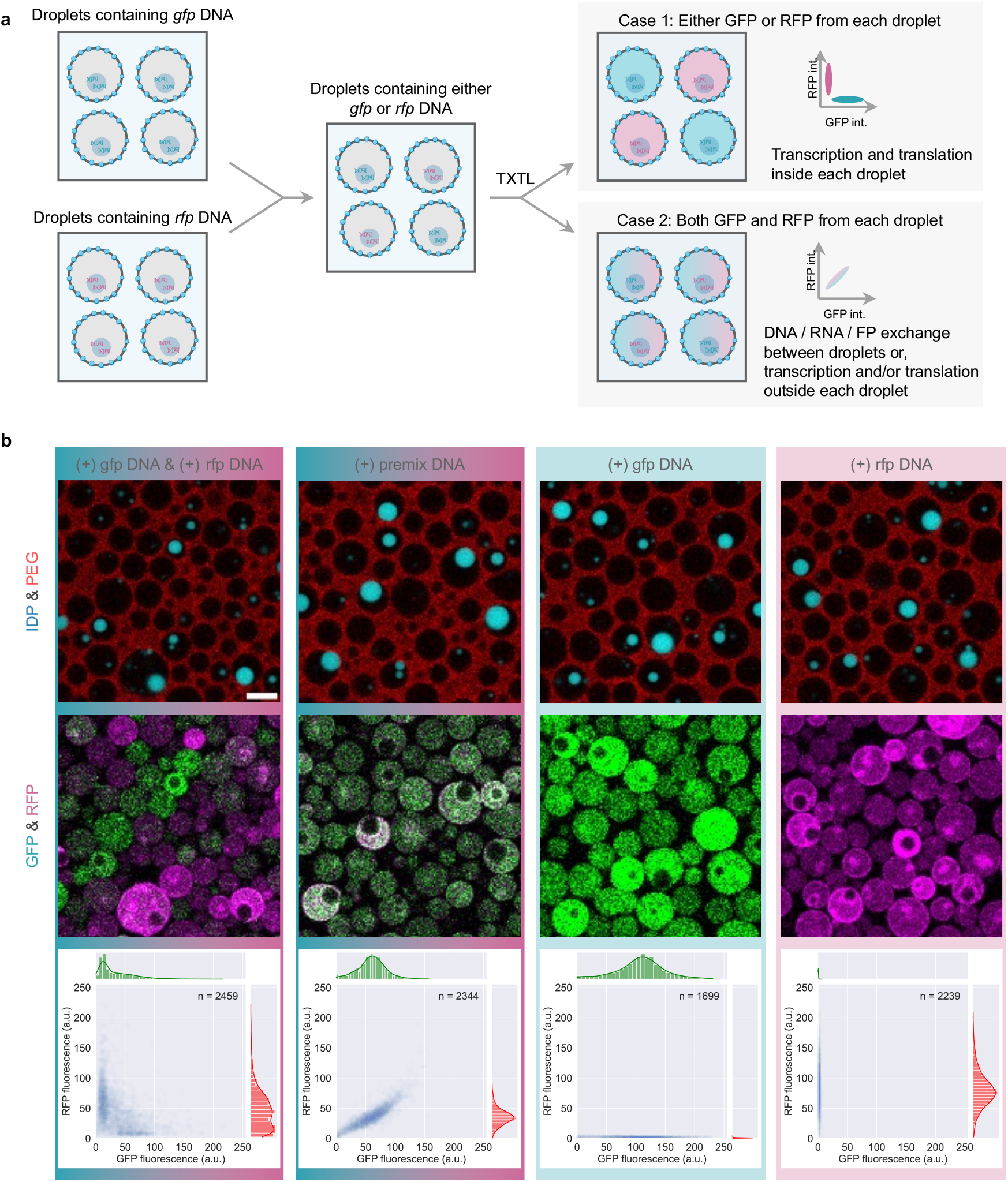
Genotype - phenotype correspondence. **a** Schematic showing how to prepare droplets containing homogeneous genotypes and inquire whether homogeneous phenotypes are expressed in each droplet. **b** Expression of GFP and RFP at 120 min from the initial image acquisition. The first column at the left end shows the sample prepared in the manner depicted in **a**. The second column shows the condition where premixed gfp DNA solution and rfp DNA solution were added. The third and fourth columns show conditions where gfp DNA and rfp DNA are solely added, respectively. The bottom row shows marginal plots, where scatter plots showing GFP and RFP fluorescence from each droplet are combined with histograms for each axis. n values indicate the number of droplets plotted for each condition. The scale bar represents 10 μm.

### Localized translation in outer Dex droplet

We aimed to conduct translation in the outer Dex droplet in our droplet-in-droplet system coupled with the transcription at the inner IDP droplet (Figure 3a). Before transcription-translation experiments, to confirm the partitioning of ribosomes, we labeled them with dye and observed their locations. Figures 3b and 3c illustrate that ribosomes are preferentially localized in the Dex phase. The calculated enrichment factor, defined as the ratio of fluorescence in the IDP droplet to that in the Dex droplet, was found to be 0.3. The exclusion of ribosomes from the IDP droplet could be attributed to their size (∼20 nm), surpassing the previously reported coacervate’s mesh size from the same IDP (RGG-rich domain from LAF-1) of 3∼8 nm^35^.

Before microscopic observations of transcription and translation, the IDP-Dex two-phase system in conventional test tubes was macroscopically separated to examine the localization of translation activity, similar to the method used for identifying transcription sites (Supplementary Figure 4a). The addition of mRNA to express GFP (mStayGold QC2-6 FIQ^36^) resulted in comparable signal increases in both the whole mixture (IDP and Dex phases) and the supernatant Dex phase, suggesting that translation preferentially proceeds at the Dex phase rather than the IDP phase (Supplementary Figure 4e). The mRNA, containing an aptamer sequence (F30-8Pepper), showed equal levels in both whole mixture and supernatant conditions, indicating that the IDP phase does not enrich mRNA and mRNA is readily available at the Dex phase for translation (Supplementary Figure 4d).

Based on these results, we aimed to couple transcription and translation in the droplet-in-droplet structure and conducted time-lapse observation under the fluorescence microscope. The protein expression was monitored by GFP fused with an N-terminal SNAP-tag designed to covalently bind with benzyl-guanine-modified Dextran (BG-Dex) so that GFP was partitioned to the Dex phase (Figure 3d). The effect of this interaction between SNAP-GFP and BG-Dex was assessed by comparing conditions with or without BG-Dex. Although the expression level of SNAP-GFP was independent of the presence of BG-Dex, as detected by endpoint measurements with a plate reader, microscopic observations revealed that SNAP-GFP was not partitioned to the Dex phase when BG-Dex was not added (Supplementary Figure 5).

Figure 3e shows that the fluorescence signal from SNAP-GFP continuously increased at the Dex phase within 240 min of observation, while no signal increase was confirmed if the template DNA was not added. Additionally, Dex droplets were distinguished by the presence or absence of inner IDP droplets. Droplets containing IDP droplets exhibited approximately twice the GFP signal compared to those without. The enhanced GFP production suggests that the IDP phase accumulates DNA molecules, enhancing the efficiency of transcription and translation.

### Cross-talk assessment among droplets

In artificial cells, it is crucial to ensure that the genotype corresponds to the phenotype and molecules that represent the phenotype are kept within each compartment. Given that the droplets used here are membrane-less and offer molecular permeability, we investigated how the coupling of genotype (i.e., DNA) with phenotype molecules (i.e., proteins to be expressed) is maintained among droplets.

We prepared two sets of droplets; one carried DNA encoding for GFP (mStayGold QC2-6 FIQ) fused with a SNAP-tag, and the second with DNA for RFP (mScarlet-I^37^) fused with a SNAP-tag. We then investigated whether GFP or RFP was expressed exclusively in each droplet after the two populations of droplets were gently mixed and added to a well for observation (Figure 4a).

We also analyzed droplets in which the premix solution of gfp DNA and rfp DNA was added before droplet formation to serve as a reference for fully fused conditions. Additional control conditions included droplets containing only a single set of either GFP DNA or RFP DNA for comparison purposes. In all conditions, the total final DNA concentration was set to 0.1 nM.

Figure 4b demonstrates that when droplets containing either gfp or rfp DNA are imaged, a distinct expression of either GFP or RFP tends to be observed at 120 min, as evidenced by the orthogonal distribution in the scatter plot. Conversely, when gfp and rfp DNAs were premixed prior to the formation of droplet-in-droplet structures, a clear correlation of GFP and RFP fluorescence among each droplet was observed. Together with conditions added with all gfp DNA or all rfp DNA, where either GFP or RFP was solely expressed, these results suggest that despite the proximity, each droplet largely retains its genotype-phenotype correspondence over the duration of protein expression.

This orthogonality was further evaluated by converting the scatter plot into polar coordinates after subtraction and normalization, focusing on the θ values for each condition (Supplementary Figure 6a). This procedure was applied across all 15 frames captured (0 to 150 min with 10 min intervals). We then calculated a symmetry index defined as |θ – 45°| and plotted it for each time point and condition (Supplementary Figure 6b). Although the symmetry index values showed variability within the initial 50 minutes for all conditions, likely due to noise from limited expression levels (r in polar coordinates), these values did not increase and rather stabilized after 50 minutes, even in the (+) gfp DNA & (+) rfp DNA condition (Supplementary Figure 6c). This suggests that the co-expression of GFP and RFP observed in a minority of droplets is due to time-independent factors, such as the initial DNA distribution in the droplet formation step, rather than time-dependent factors, such as transcription, translation, or the diffusion of their products (mRNA and fluorescent proteins).

## Discussion

Our study demonstrated that (1) droplet-in-droplet structures can be engineered in an all-aqueous system by combining a coacervate of intrinsically disordered proteins (IDPs) with interface-stabilized Dex-PEG aqueous two-phase systems (ATPS); (2) transcription can be actively carried out within the inner IDP droplet, which acts as a “nucleus”; and (3) translation in the outer Dex droplet can be coupled to transcription, mimicking the “cytosol.” Crucially, Dex droplets containing inner IDP droplets exhibited enhanced protein expression compared to those lacking IDP droplets. As evidenced by the specific protein phenotypes dictated by the DNA encapsulated within the inner IDP droplets, both transcription and translation processes were carried out independently in each droplet. This observation confirms that our membrane-free artificial cell system effectively maintains a correspondence between genotype and phenotype, preserving the individuality of each functional compartment.

The membrane-less nature of the inner IDP coacervate facilitates the seamless transfer of transcribed mRNA to the outer droplet, triggering translation. This eliminates the need for complex molecular systems such as pores or transporters often required by enclosed compartments like liposomes or proteinosomes. The inner IDP droplet was designed to enrich enzymes (RNA polymerase) and template DNAs through the genetic integration of peptide interaction modules. Given the established utility of other orthogonal interacting peptide pairs^32^ and recent advancements in the mechanism of the coexistence of distinct membrane-less organelles^38,39^, the introduction of additional modular artificial organelles appears feasible.

The outer Dex droplets, surrounded by colloidal emulsifiers rather than layers of membranes, maintain molecular permeability inherent to ATPS while preventing coalescence. This characteristic will be critical for enabling a sustained energy supply within the artificial cell system, which could promote chemical perturbations and the production of biologically active molecules.

Furthermore, the spatial segregation of transcription and translation within our system sets the stage for spatiotemporal control over these processes. For example, if transcription and translation lead to the production of repressors accumulated within the inner droplet, a negative feedback mechanism could be triggered. Adjusting the balance between repression and diffusion of these repressors could induce oscillatory behaviors in transcription and translation activities akin to the cellular cycle. The inherent permeability of the Dex droplets further enhances this model by allowing continuous resource supplementation, thereby supporting long-term observations to increase the utility of this artificial cell model for experimental and practical applications.

## Method

### Plasmid DNA and linear template DNAs

Plasmids and linear template DNAs used in this study are listed in Supplementary Information. Their sequences were all verified by Sanger sequencing. Plasmid DNA was purified by FastGene PlasmidMini (NIPPON Genetics, FG-90402). Linear DNA was amplified by PCR (KOD One PCR Master Mix, TOYOBO, KMM-101) and purified (FastGene Gel / PCR Extraction Kit, NIPPON Genetics, FG-91202).

### Expression and purification of recombinant proteins

IDP (SZ1-RGG-BFP-RGG) was cloned in a pRSET-B vector with an N-terminal His-rich tag, expressed in T7 Express *lysY/Iq* Competent *E. coli* (NEB, C3013I) and purified as previously reported with some modifications^25^. Fresh colonies on LB agar plate with Carbenicillin (100 μg/ml) were directly inoculated to 1 L of LB liquid media containing the same concentration of Carbenicillin. The culture was incubated at 37°C with shaking at 200 rpm until the OD_600_ reached 0.8 - 1.0 and induced by 1 mM of isopropyl β-D-1-thiogalactopyranoside (IPTG) overnight by shaking at 200 rpm at 18°C. Cells were then harvested by centrifugation at 8000 rpm for 15 min at 4°C and resuspended by 50 ml of lysis buffer (20 mM Tris-HCl pH=7.5, 1 M NaCl, 20 mM Imidazole). Cells were disrupted by sonication and incubated in a 60°C water bath for 1 - 2 h so that contaminating proteins denature and aggregate. This solution was centrifuged at 15,000 g for 1 min, and the supernatant was centrifuged at the same condition again. The second supernatant was loaded onto Ni Sepharose 6 Fast Flow (Cytiva) equilibrated with 2CV of lysis buffer at room temperature. The column was washed with over 2CV of wash buffer (20 mM Tris-HCl pH=7.5, 0.5 M NaCl, 20 mM Imidazole), and the IDP was eluted with elution buffer (20 mM Tris-HCl pH=7.5, 0.5 M NaCl, 500 mM Imidazole). Fractions of the eluted solution were immediately moved to the 60°C water bath so as not for IDP to phase separate at room temperature. 3 mL of the purified IDP at 60°C was loaded onto Econo-Pac 10DG Desalting Prepacked Gravity Flow Column (BIO-RAD) equilibrated with 20 mL of stock buffer (50 mM HEPES-KOH pH=7.5, 0.5 M NaCl) in an incubator set to 60°C. The desalted IDP solution was eluted by 4 mL of the stock buffer and moved back to the 60°C water bath. This stock solution was aliquoted to PCR tubes on a heat block set to 60°C, flash-frozen with liquid nitrogen, and stored at -80°C.

SZ2-T7RNAP was cloned in the pBAD vector with an N-terminal His-rich tag and expressed from One Shot TOP10 Chemically Competent (Invitrogen C404003) and purified. A single fresh colony on an LB agar plate with Carbenicillin (100 μg/ml) was inoculated to 5 mL of LB liquid media containing Carbenicillin (100 μg/ml) for pre-culture.

This pre-culture was transferred to 1 L of LB liquid media containing Carbenicillin (100 μg/ml). The culture was shaken at 200 rpm at 37°C until OD_600_ reached 0.6 and induced by 0.02% L-arabinose (stock solution: 20%) overnight by shaken at 200 rpm at 18°C. The cells were harvested by centrifugation at 8000 rpm for 15 min at 4°C and resuspended by 50 ml of wash buffer (20 mM Tris-HCl pH=7.5, 0.5 M NaCl, 20 mM Imidazole, 5 mM 2-mercaptoethanol). Cells were disrupted by sonication and ultracentrifuged (81,000 g, 4°C, 20 min). The supernatant was loaded onto Ni Sepharose 6 Fast Flow (Cytiva) equilibrated with 2CV of wash buffer. The following operations were conducted at 4°C unless otherwise specified. The column was washed with over 2CV of wash buffer, and the protein was eluted with elution buffer (20 mM Tris-HCl pH=7.5, 0.5 M NaCl, 250 mM Imidazole, 5 mM 2-mercaptoethanol). Fractions of the eluted solution were concentrated by Amicon Ultra Centrifugal Filter, 50 kDa MWCO. This solution was buffer exchanged by Econo-Pac 10DG Desalting Prepacked Gravity Flow Column (BIO-RAD) equilibrated with 20 mL of 2× stock buffer (100 mM HEPES-KOH pH=7.5, 0.2 M NaCl, 40 mM 2-mercaptoethanol, 2 mM EDTA-NaOH pH=8.0, 2 mM DTT) at 4°C. Glycerol was added to the eluted solution as stored at -20°C. T7RNAP without an SZ2 tag was prepared in almost the same manner, with the exception that this protein was cloned in the pQE30 vector and induced by 1 mM of isopropyl β-D-1-thiogalactopyranoside (IPTG) at OD_600_ = 0.8 ∼ 1.0.

SZ2-LacR was cloned in the pBAD vector with a C-terminal 6×His tag. By replacing the C-terminal region of the original LacR with His tag, this construct is not supposed to form a tetramer (dimerized dimers) but to form a dimer, which is reported to be sufficient for binding lacO DNA. This construct was expressed from One Shot TOP10 Chemically Competent (Invitrogen C404003) and purified. A single fresh colony on an LB agar plate with Carbenicillin (100 μg/ml) was inoculated to 5 mL of LB liquid media containing Carbenicillin (100 μg/ml) for pre-culture. This pre-culture was transferred to 1 L of LB liquid media containing Carbenicillin (100 μg/ml). The culture was shaken at 200 rpm at 37°C until OD_600_ reached 0.5 and induced by 0.02% L-arabinose (stock solution: 20%) for four hours by shaken at 200 rpm at 30°C. The cells were harvested by centrifugation at 8000 rpm for 15 min at 4°C and resuspended by 50 ml of lysis buffer (20 mM Tris-HCl pH=7.5, 1 M NaCl, 20 mM Imidazole). Cells were disrupted by sonication and ultracentrifuged (81,000 g, 4°C, 20 min). The supernatant was loaded onto Ni Sepharose 6 Fast Flow (Cytiva) equilibrated with 2CV of wash buffer (20 mM Tris-HCl pH=7.5, 0.5 M NaCl, 20 mM Imidazole). The following operations were conducted at 4°C unless otherwise specified. The column was washed with over 2CV of wash buffer, and the protein was eluted with elution buffer (20 mM Tris-HCl pH=7.5, 0.5 M NaCl, 500 mM Imidazole). Fractions of the eluted solution were concentrated by Amicon Ultra Centrifugal Filter, 10 kDa MWCO, and applied to a size exclusion column, Superdex 200, 10/200GL (Cytiva, 17-5175-01) equilibrated with stock buffer (200 mM Tris-HCl pH=7.5, 200 mM KCl, 1 mM EDTA, 1 mM DTT). Fractions within the single peak were collected and concentrated again by Amicon Ultra Centrifugal Filter, 10 kDa MWCO. This stock solution was aliquoted to PCR tubes on ice, flash-frozen with liquid nitrogen, and stored at -80°C.

### Preparation of BLG colloids and their PEGylation

Colloidal microgels of BLG (beta-lactoglobulin) were prepared according to Gonzalez-Jordan, A. *et al*^30^. The solution of 4.4 mM CaCl_2_ and 40 mg/ml BLG (Sigma-Aldrich, L3908) dissolved in Milli-Q water was incubated at 80°C for 15 hours, during which the solution turned to white turbid to form BLG colloids. For PEGylation, this BLG colloid solution (10 mg/ml) was mixed with 16.3 mg/ml mPEG2k-NHS (NANOCS, PG1-SSA-2k) freshly dissolved in DMSO (initial concentration: 81.5 mg/ml) and 50 mM HEPES-KOH pH=8.3 and incubated for 1.5 hours. After that, the solution was washed with 40 mM HEPES-KOH pH=7.6 by Amicon Ultra Centrifugal Filter, 100 kDa MWCO repeatedly for four times. Finally, the same buffer was added until the volume reached a quarter of the initial volume to make BLG concentration back to 40 mg/ml and stored at 4°C. The turbidity was measured at OD_700_ by NanoDrop (Thermo Scientific) to be 0.2 at 1 mm of optical path length.

### Preparation of Atto647N-PEG

Atto647N dye-labeled PEG was prepared from PEG20k-diamine (Sigma-Aldrich, 14509) and Atto647N-NHS (Sigma-Aldrich, 18373). 100 mM HEPES-KOH pH=8.3, 1 wt% PEG20k-diamine, 1.26 g/l Atto647N-NHS dissolved in DMSO (initial concentration: 10 g/l) were mixed with Milli-Q water and incubated for 1.5 hours. Unreacted dyes were excluded by Zeba Spin Desalting Column (Thermo Scientific, 89882) according to the manufacturer’s instruction. The product was diluted to 0.1 wt% by Milli-Q water and stored at 4°C.

### Preparation of BG-Dex

Benzyl-guanine conjugated Dextran was prepared from Amino Dextran 500k (Invitrogen, D7144) and BG-GLA-NHS (NEB, S9151S). 50 mM HEPES-KOH pH=8.3, 1 wt% Amino Dextran, 0.16 mM BG-GLA-NHS freshly dissolved in DMSO (initial concentration: 10 g/l) were mixed with Milli-Q water and incubated for 1.5 hours. Unreacted BG-GLA-NHS was washed out with Milli-Q water by Amicon Ultra Centrifugal Filter, 100 kDa MWCO repeatedly for four times. Finally, Milli-Q water was added until the volume reached two-thirds of the initial volume to make the conjugated Dextran concentration 1.5 wt% and stored at 4°C.

### Labeling lacO DNA, T7RNAP, and ribosomes with FL (fluorescein)

FL-labeled lacO DNA was prepared by PCR labeling with FL-dUTP (Roche, 11373242910). PCR enzyme and 10x PCR buffer are from TaKaRa Taq Hot Start Version (Takara, R007A). The PCR mixture was composed of 1x PCR buffer, 0.05 U/μL of PCR enzyme, 0.1 mM each of dATP, dCTP, dGTP, 0.05 mM of dTTP, 0.05 mM of FL-dUTP, 0.3 μM each of forward and reverse primers, and 20 pg/μl of template DNA. The amplicon was designed to contain a lacO sequence. The details about sequences of primers and the amplicon can be found in supplementary information. The PCR product was purified by FastGene Gel / PCR Extraction Kit (NIPPON Genetics, FG-91202) according to the manufacturer’s instructions. The degree of labeling was calculated to be 356 dyes per dsDNA.

FL-labeled SZ2-T7RNAP and T7RNAP were prepared from purified SZ2-T7RNAP, T7RNAP (mentioned above), and FL-NHS (Thermo Scientific, 46410). 10 g/l SZ2-T7RNAP or T7RNAP, 0.7 g/l FL-NHS freshly dissolved in DMSO (initial concentration: 10 g/l) were mixed with labeling buffer (50 mM HEPES-KOH pH=8.3, 500 mM NaCl dissolved in Milli-Q water). The mixture was incubated at 4°C for two hours. Unreacted dyes were excluded by exchanging the buffer to stock buffer (20 mM HEPES-KOH pH=7.6, 100 mM Potassium acetate, 7 mM 2-mercaptoethanol dissolved in Milli-Q water) by Zeba Spin Desalting Column (Thermo Scientific, 89882) according to the manufacturer’s instruction. The degrees of labeling were calculated to be 0.35 dye per SZ2-T7RNAP and 0.24 dye per T7RNAP.

FL-labeled ribosomes were prepared from ribosome solution (Solution III of PURE frex2.0, GeneFrontier) and FL-NHS (Thermo Scientific, 46410). 4.0 μM ribosomes, 0.03 g/l FL-NHS freshly dissolved in DMSO (initial concentration: 10 g/l) were mixed with labeling buffer (50 mM HEPES-KOH pH=8.3, 10 mM Magnesium acetate, 100 mM KCl dissolved in Milli-Q water). The mixture was incubated at 4°C for two hours. Unreacted dyes were excluded by exchanging the buffer to stock buffer (40 mM HEPES-KOH pH=7.6, 10 mM Magnesium acetate, 30 mM KCl, 7 mM 2-mercaptoethanol dissolved in Milli-Q water) by Zeba Spin Desalting Column (Thermo Scientific, 89882) according to the manufacturer’s instruction. The degree of labeling was calculated to be 0.3 dye per ribosome.

### Preparation of 384-well Microscopy Plates

384-well glass bottom microplate (Corning, 4581) was added with 25 μL per well of 5% Cell Cleaning Solution (SANSYO, T-A-28), incubated for 4h at 37°C and rinsed copiously with Milli-Q water. Silica was etched with 25 μL per well of 8N KOH for 1h at room temperature and rinsed copiously with Milli-Q water, followed by ethanol. Silanization was conducted overnight by adding 25 μL per well of 2% Dichlorodimethylsilane (Sigma, 440272) dissolved in a solvent of 95% ethanol, 4.99% Milli-Q water, and 0.01% acetic acid, and rinsed copiously with ethanol. The plate was sealed with an adhesive PCR foil. Just before use, silanized wells were opened by cutting the foil, washed by pipetting 20 μL per well of Milli-Q water, and 20 μL per well of 5% Pluronic F-127 was added, and incubated for 5min. Then, the remaining solution was sucked up and washed again by pipetting 50 μL per well of Milli-Q water. Sample solutions were immediately added to the vacant wells.

### Microscopy and image analysis

All images were acquired by TCS-SP8 X (Leica Microsystems) confocal laser scanning microscope equipped with white-light laser lines and HyD detectors. Acquired images were cropped and/or merged with Fiji and analyzed using custom Python codes available here: https://github.com/tomohara97/droplet-in-droplet. Microscopy configurations can be found in the supplementary information.

### Sample preparations for microscopy observations

For the observation of the droplet-in-droplet structure with four channels (IDP, Dex, PEG, colloidal emulsifiers), the components were mixed as follows: IDP (2.4 μM), Dex 500k (1 wt%), TRITC-Dex (0.01 wt%), mPEG2k-BLG (2.4 mg/ml), Atto647N-PEG (0.001 wt%), SYPRO orange in DMSO (10x), and PEG 35k (8 wt%) were sequentially added. For other observations, samples contained the following components unless otherwise noted: IDP (3.2 μM), SZ2-LacR (1 μM), PEG 35k (6 wt%), PURE frex 2.0 Solution 1 (0.54x), PURE frex 2.0 Solution 2 ΔT7RNAP (1x), PURE frex 2.0 Solution 3 (1x), mPEG2k-BLG (2.4 mg/ml), Atto647N-PEG (0.001 wt%), Dex 500k (0.1 wt%), SZ2-T7RNAP (20 nM). Some components were mixed beforehand as a master mix, which typically included PURE components (Solution 1∼3), mPEG2k-BLG, Atto647N-PEG, Dex 500k, and SZ2-T7RNAP. All PURE frex 2.0 components were purchased from GeneFrontier, and PURE frex 2.0 Solution 2 ΔT7RNAP was custom-ordered. Additional specific information for each observation is specified below.

To observe and compare FL-labeled T7RNAP, not every experimental condition included SZ2-T7RNAP and SZ2-LacR. Instead, one of the following was added for comparison: 1 μM of labeled T7RNAP, 1 μM of unlabeled T7RNAP, an optically equivalent concentration of FL, or Milli-Q water.

To observe and compare the interaction between FL-labeled lacO DNA and SZ2-LacR, 0.1 nM of FL-labeled lacO DNA was added with or without SZ2-LacR (1 μM).

To observe transcription activities, 3 μM of HBC530 (a kind gift from Yuki Goto, Kazuhito V. Tabata, and Mana Ono, synthesized and purified according to the previous report^33^, initially dissolved in DMSO as 300 μM stock) was added with or without linear template DNA. This template DNA encodes the F30-8Pepper gene under the control of the T7 promoter, followed by the T7 terminator and two repeats of the lacO sequence. To observe FL-labeled ribosomes, 0.1 μM of labeled ribosomes or an optically equivalent concentration of FL was added.

To observe the expression of GFP-SNAP and RFP-SNAP, template DNA at a concentration of 0.1 nM was supplemented with 0.015 wt% benzyl-guanine conjugated Dextran (BG-Dex). To verify genotype-phenotype correspondence, three distinct droplet-in-droplet structures were prepared by varying the DNA added: (1) GFP-SNAP coding DNA, (2) RFP-SNAP coding DNA, and (3) a premixed solution containing equal parts of (1) and (2). Each mixture was vigorously agitated to ensure uniformity. Subsequently, equal volumes of the mixtures from (1) and (2) were combined gently in a tube and then carefully transferred to a 384-well plate without any agitation.

All samples were vigorously agitated in the tube unless otherwise specified and loaded onto a 384-well microscopy plate. The plate was mildly centrifuged at 700 rpm for 1 min to maximize the number of droplets to be observed near the glass surface at the bottom. Wells with samples were sealed by adding 10 μL of mineral oil (Sigma-Aldrich, M5904) and subjected to microscopy observation.

## Supporting information

Supplemental informatoin

DNA information

Microscopy information

## Acknowledgments

We thank members of the Noji lab for valuable discussions and critical assessment of the work. We are also grateful to Yuki Goto, Kazuhito V. Tabata, and Mana Ono for generously providing their reagents. This work was supported in part by JST CREST, Japan (JPMJCR19S4), to H.N., Grants-in-Aid for Scientific Research (S), JP19H05624 to H.N. from the Japan Society for the Promotion of Science.

## Author Contributions

K.T., Y.M., and H.N. designed the research. K.T. performed experiments and analyses. K.T. and H.N. wrote the manuscript.

