## Supplemental informatoin for "Artificial cells with all-aqueous droplet-in-droplet structures for spatially separated transcription and translation"

### Supplementary Methods

#### in vitro transcription

RNA was synthesized by *in vitro* Transcription T7 Kit (Takara, 6140) and purified by NucleoSpin RNA Clean-up XS (MACHEREY-NAGEL, 740903.50) according to manufacturers' instructions.

#### Endpoint fluorescence measurement by a plate reader

For endpoint fluorescence measurement after real-time imaging, the 384-well plate was transferred to SpectraMax iD3 (Molecular Devices), and wells were read for each fluorescence with the following settings: excitation at 500 nm, detection at 540 nm, read from the bottom, temperature control OFF (25.5°C).

#### Real-time fluorescence measurement by qPCR machine

For real-time fluorescence measurement of transcription and translation in reconstituted IDP-Dex systems, sampled mixtures were prepared as quadruplicate, added to 0.2 ml 8-Tube PCR Strips without Caps (Bio-Rad, TLS0851) and sealed with 0.2 ml Flat PCR Tube 8-Cap Strips (Bio-Rad, TCS0803). Tubes were set in CFX Opus 96 Real-Time PCR Instrument (Bio-Rad, 12011319J1). Measurement was conducted in two channels (red "ROX" channel for HBC620 activation representing transcription and green "FAM" channel for GFP expression representing translation), 10 min intervals, and 30°C (isothermal).

#### Sample preparations for microscopy observations

To evaluate the effect of Dex-PEG interface stabilization facilitated by PEG-BLG (Supplementary Figure 1), the following components were pre-mixed as a master mix: Tris-HCl at pH 7.5 (14 mM), Dextran 550k (1.95 wt%), TRITC-Dextran 500k (0.05%), and Thioflavin T (20 µM). Following distribution of this master mix, PEG-BLG colloids, or BLG colloids (1 g/L), were introduced along with PEG 35k (8 wt%).

For examining the partitioning of SNAP-GFP into the Dex phase mediated by BG-Dex (Supplementary Figure 5), the following components were included: IDP (3.2 µM), SZ2-LacR (1 µM), PEG 35k (6 wt%), PURE frex 2.0 Solution 1 (0.54x), PURE frex 2.0 Solution 2 ΔT7RNAP (1x), PURE frex 2.0 Solution 3 (1x), mPEG2k-BLG (2.4 mg/ml), Atto647N-PEG (0.001 wt%), Dex 500k (0.1 wt%), and SZ2-T7RNAP (20 nM). A master mix was prepared in advance, comprising PURE components (Solution 1~3), mPEG2k-BLG, Atto647N-PEG, Dex 500k, and SZ2-T7RNAP. Depending on the conditions, 0.015 wt% of BG-Dex and template DNA (0.1 nM) were added.

**Supplementary Figure 1 | Stabilization of the Dex-PEG Interface via proteinaceous colloids.**

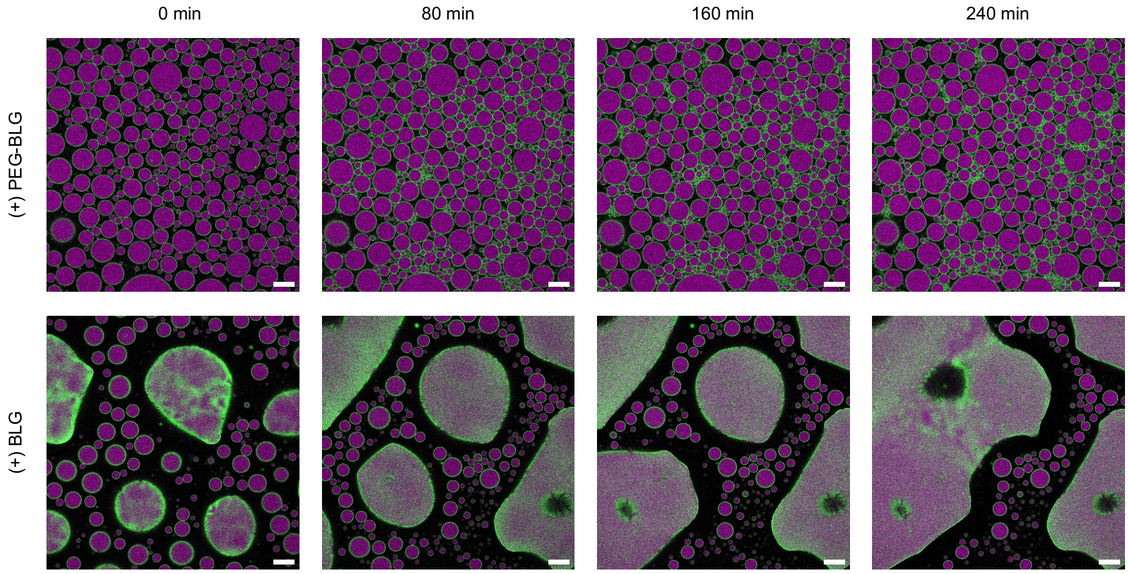

**Supplementary Figure 1 | Stabilization of the Dex-PEG Interface via proteinaceous colloids.** Either PEGylated beta-lactoglobulin (BLG) microgels (PEG-BLG) or BLG microgels were added to a Dex-PEG aqueous two-phase system (ATPS). The system includes Dextran 550k (1.95 wt%), TRITC-Dextran 500k (0.05%), PEG 35k (8 wt%), Thioflavin T (20 μM) and proteinaceous colloids (1 g/L), buffered with Tris-HCl at pH 7.5 (14 mM). The colloids are visualized using Thioflavin T (green), while the Dex phase is labeled with TRITC-Dex (magenta). All scale bars represent 50 μm.

**Supplementary Figure 2 | Enrichment of RNAP into the IDP droplet with or without SZ2 tag.**

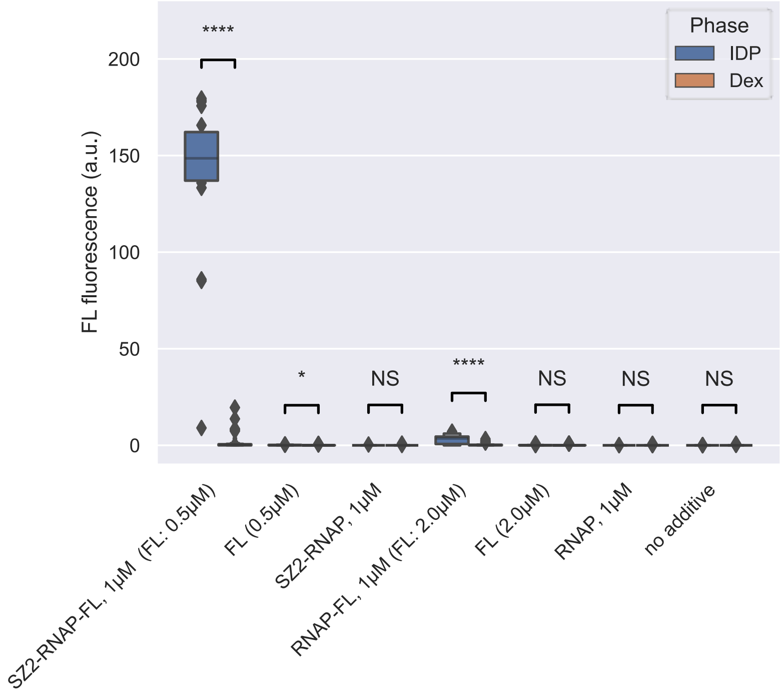

**Supplementary Figure 2 | Enrichment of RNAP into the IDP droplet with or without SZ2 tag.** Extended data from Figure 2b. The droplet-in-droplet system was added with one of the following for comparison: 1 μM of labeled RNAP, 1 μM of unlabeled RNAP, an optically equivalent concentration of FL, or Milli-Q water. Significance was assessed using a two-tailed Welch's t-test: NS ($p > 0.05$), * ($p < 0.05$), ** ($p < 0.01$), *** ($p < 0.001$), **** ($p < 0.0001$).

**Supplementary Figure 3 | Location of RNA expressed in the droplet-in-droplet structure.**

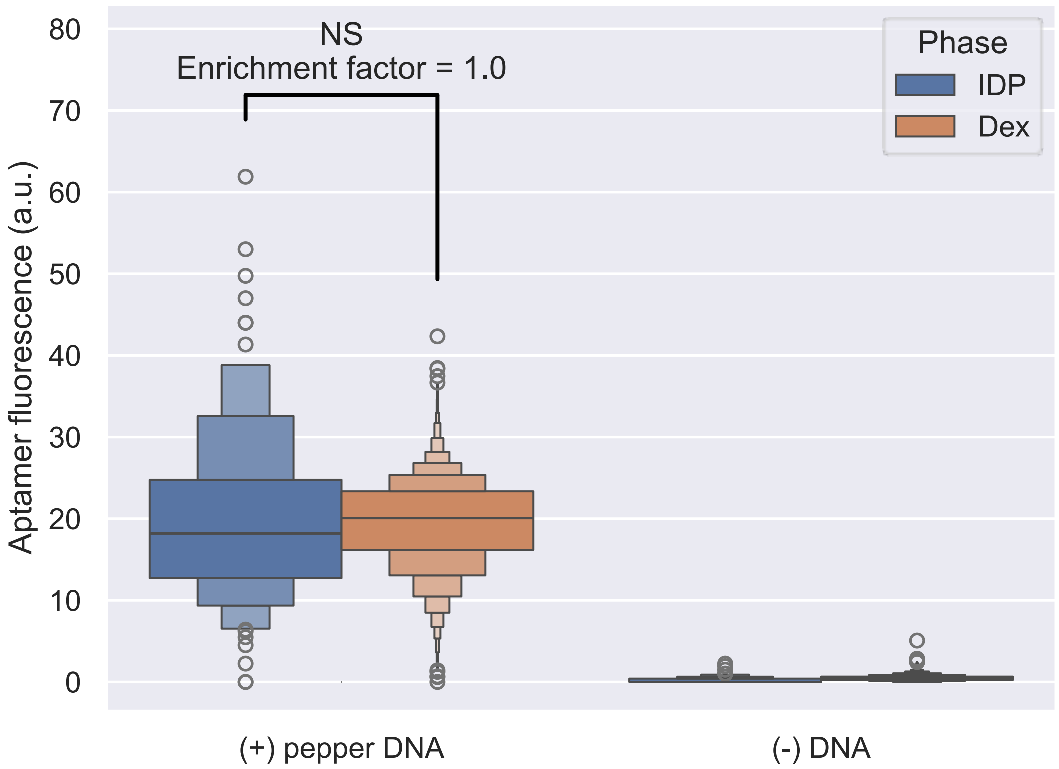

Supplementary Figure 3 | Location of RNA expressed in the droplet-in-droplet structure.

Extended data from Figure 2g, showing the distribution of aptamer fluorescence after 480 minutes of observation. The data compare conditions with and without DNA in either the inner IDP droplet or the outer Dex droplet. Statistical significance was assessed using a two-tailed Welch's t-test, with the following notation: NS (not significant, $p > 0.05$), * ($p < 0.05$), ** ($p < 0.01$), *** ($p < 0.001$), **** ($p < 0.0001$).

**Supplementary Figure 4 | Transcription and translation in reconstituted IDP-Dex two-phase system.**

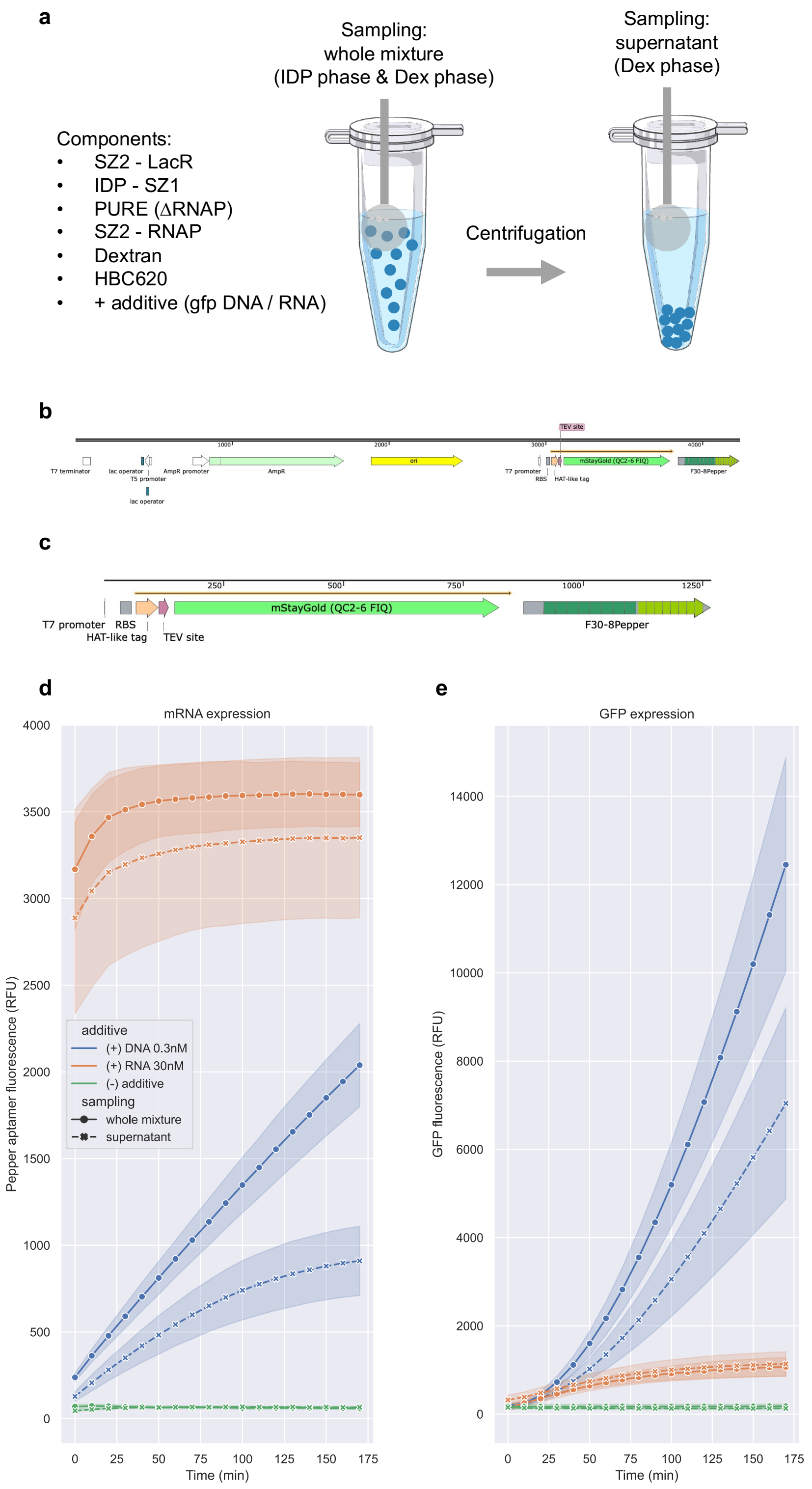

**Supplementary Figure 4 | Transcription and translation in reconstituted IDP-Dex two-phase system.** The transcription and translation activities within the reconstituted IDP-Dex two-phase system. **a** The upper Dex phase was extracted after centrifugation and compared with the pre-centrifugation sample containing the IDP phase. Both samples underwent real-time monitoring of transcription and translation activities. Transcription was tracked using the pepper aptamer and HBC620 fluorescence (red channel), while translation was observed through GFP expression (green channel). **b** The template DNA for monitoring both transcription and translation included two lacO sequences and the GFP and pepper genes under the T7 promoter. **c** For translation monitoring, the template RNA was synthesized in vitro from the DNA outlined in **b**, followed by membrane-column purification. **d** Real-time monitoring of transcription via the red channel (HBC620 activation by pepper RNA expression) and **e** translation through the green channel (GFP expression) under one of three conditions: 0.3 nM template DNA, 30 nM template RNA, or no additives. The experiments were conducted in quadruplicate, and the results are presented as means with error bands representing 2 SE.

**Supplementary Figure 5 | Partitioning expressed GFP-SNAP within the Dex droplet by BG-Dex.**

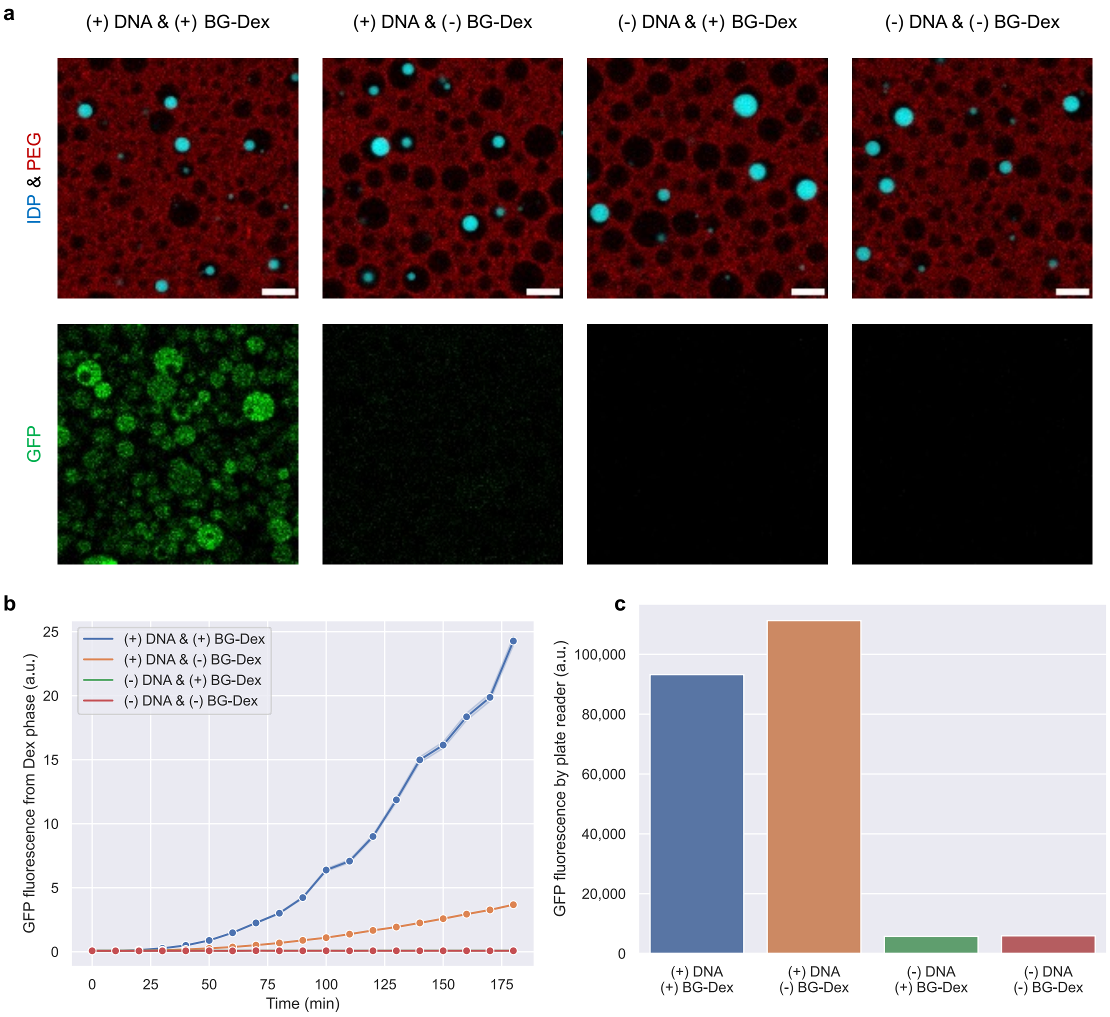

**Supplementary Figure 5 | Partitioning expressed GFP-SNAP within the Dex droplet by BG-Dex.** The droplet-in-droplet structures were formed with or without BG-Dex (0.015%) and with or without template DNA coding GFP-SNAP (0.1 nM). **a** Locations of expressed GFP-SNAP after 180 min of confocal imaging. Scale bars indicate 10 µm. **b** Time trajectories of GFP fluorescence within the Dex phase. Error bands represent 2SE. **c** The endpoint measurement of GFP fluorescence from the same samples, taken immediately following the 180 min imaging using a plate reader.

### Supplementary Figure 6 | Analysis of the orthogonality in GFP / RFP expression

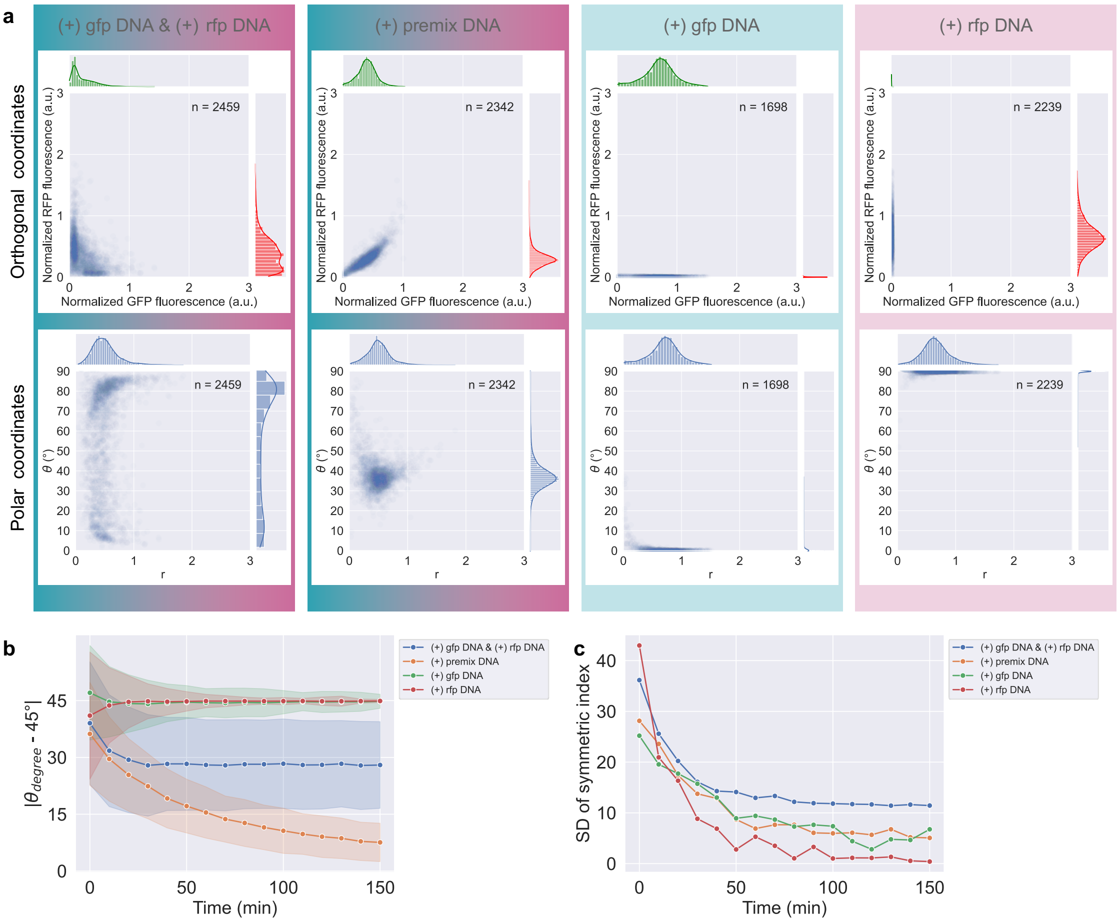

**Supplementary Figure 6 | Analysis of the orthogonality in GFP / RFP expression.** Extended data from Figure 4b. **a** In the upper row, data at 120 min shown in Figure 4b were subtracted and normalized. The optical backgrounds for the GFP and RFP channels were defined using a sample with no DNA added and applied for subtraction. Mean intensity values for GFP and RFP in the (+) gfp DNA and (+) rfp DNA conditions, respectively, at the final frame (150 min), were used for normalization. In the lower row, the orthogonal coordinates from the upper row were transformed into polar coordinates. Data points were filtered by a threshold of r (expression level), defined as the 95% quantile in the condition where DNA was not added. **b** The above-mentioned analysis was applied across all 15 frames captured (0 to 150 min), and the time trajectories of the symmetric index defined as |θ – 45°| were shown. Error bands represent 1 SD. **c** The SD values in **b** were plotted for each frame and each condition.
